# Spatial Recruitment Model for Motor Potentials Evoked By Transcranial Magnetic Stimulation (TMS) for In-Silico Testing and Tuning of Closed-Loop Procedures

**DOI:** 10.64898/2026.06.25.734597

**Authors:** Maryam Farahmandrad, Mahdi Bayati, Victoria Middleton, Jennifer Bowman, Nancy Donachie, Joseph Kriske, John Kriske, Jonathan Downar, Stefan Goetz

## Abstract

The primary motor cortex has a prominent role in transcranial magnetic stimulation (TMS). It is one of few locations that provide directly observable responses, and its physiology serves as model or reference for almost all other TMS targets, e.g., through the motor threshold and spatial targeting relative to its position. It furthermore sets the safety limits for the entire brain. Its easily detectable responses have led to closed-loop methods for a range of aspects, e.g., for automated thresholding, amplitude tracking, and targeting. The high variability of brain stimulation methods would substantially benefit from fast unbiased closed-loop methods. However, the development of more potent methods would early on in the design phase require proper models that allowed tuning and testing with sufficient conditions but without a high number of experiments, which are time-consuming and expensive or even impossible at the needed scale. On the one hand, theoretical researchers without access to experiments miss realistic spatial response models of brain stimulation to develop better methods. On the other hand, subjects should potentially not be exposed to early closed-loop-methods without sufficient prior testing. After all, initial, poorly tuned feed-back as needed for closed-loop operation is known to demonstrate erratic behavior.

To bridge this gap, we developed a digital-twin-style population model that generates motor evoked potentials in response to virtual stimuli and includes statistical information on spatial (coil position and orientation) as well as pulse-strength-dependent neural recruitment in the population to represent inter- and intra-individual variability. The population data were extracted from a combination of datasets of healthy and depressed subjects to reach large numbers and wide coverage. The model allows users to simulate different subjects and millions of runs for software-in-the loop testing. The model provides all code open-source to stimulate further development.

## I. Introduction

Transcranial magnetic stimulation (TMS) is a noninvasive method to probe and modulate brain circuits [1]. Due to easily accessible and detectable motor responses, the primary motor cortex is one of the most significant targets for brain stimulation. Stimulation of the motor cortex with enough strength can generate motor evoked potentials (MEPs), which can be recorded as a short electric wave through electromyography at the corresponding peripheral muscle. MEPs have a wide dynamic range from microvolts to millivolts [2]–[5]. Noninvasive stimulation of the primary motor cortex serves for the diagnosis and localization of motor lesions [6]. Moreover, the primary motor cortex is a preferred model for studying the neurophysiology, biophysics of brain stimulation, and development of novel technology [7]–[9] Most importantly, the motor cortex is a model circuit for most other targets for safety as well as amplitude individualization through the motor threshold [10].

MEPs in response to any brain stimulation method depend on a variety of stimulus as well as position parameters and are highly variable between subjects, within subjects from session to session, and from pulse to pulse [11]–[22] One of the most important stimulation parameters with modern focal figure-of-eight coils is accurate targeting, coil placement, and maintenance over time [23]–[29] In addition to coil location [3], [18], [30], [31], orientation [32], [33], and alignment [34], [35], the distance between the coil and the scalp [32] determine the neuronal response to TMS.

Many metrics and procedures depend on MEPs and are hampered by their complex formation and variability. Previous manual methods are increasingly automated, such as the detection of target hot-spots, motor threshold, motor excitability, and neural recruitment input–output curves [36]–[44] These methods should ideally process MEP responses in real time and adaptively adjust parameters based on previous outcomes, such as determining the next level of stimulation strength for optimal data collection with as few stimuli as possible [45], [46].

Closed-loop methods, which automatically tune one or several of the many parameters of a TMS procedure, have moved into the focus of research as they may control the high variability and remove subjectiveness as well as operator influence [47]–[49] Furthermore, automated methods can rationalize slow and labor-intensive procedures [50]–[54] Finally, closed-loop operation may enable previously not possible paradigms [55], [56].

The high nongaussian variability requires proper mathematical and statistical methods to avoid bias. The development and evaluation of such closed-loop methods typically bases on intensive testing under realistic conditions. It has been shown that widely used methods can be inaccurate and even systematically biased [1], [38], [40], [57]. For closed-loop methods, old, offline recorded experimental data are often not sufficient for testing as they do not allow sequential adaptive adjustment of stimulation parameters (coil position, stimulation [v1]strength, etc.) but are limited to the available data points. Furthermore, the development of such methods typically requires many participants, test conditions, and repeated trials already during the design phase, typically above a thousand [39], [40], [58], [59]. Such large experiments are usually not possible in an experimental study, especially in the early development stages of a method.

Closed-loop methods that only require recruitment, such as thresholding and input–output curve detection could already use available models, which stimulated the development of systematic techniques in both [44], [54], [60]–[64]. However, a model with spatial effects are missing and related closed-loop methods are highly underdeveloped and often ad-hoc. Hotspot search is for instance part of practically any TMS procedure. However, there are only few systematic procedures in the literature with little to no quantitative testing, and the majority of papers abstains from describing how the motor hotspot is found. This situation can be a problem for reproducibility and also clinical efficacy.

This article developed and provides a TMS model that generates MEP data in response to stimuli with specific stimulation strength, coil position, and coil orientation. The model is trained to population and subject statistics to generate virtual subjects that reflect typical shapes and behavior found in the population. The model incorporates inter-individual statistics of spatial hotspot location and coil orientation variability as well as recruitment and intra-individual trial-to-trial variability. As such, it is designed to confront methods during development with a high number of subjects as well as cases and can acquire large numbers of samples per subjects. Such large-scale testing with many conditions is are particularly necessary during early development phases of methods when convergence is poor or slow. The model defines virtual subjects from an entire population with their inter-individual variability, each of which further demonstrates intra-individual variability to represent a wide range of physiological responses as known from real-world settings.

The resulting framework is intended for software-in-the-loop testing of hotspot search, threshold estimation, and related adaptive TMS methods under realistic intra- and inter-individual virtual-subject variability.

## II. Methodology

### A. Clinical Data and Model Structure

We collected data from previous studies, internal databases, and the literature recorded with a range of devices (Magventure, Magstim, Rogue Research) under neuronavigation on various hand muscles (first dorsal interosseus, FDI; abductor digiti minimi, ADM; abductor pollicis brevis, APB) to cover substantial diversity of conditions and catch a wide range of properties [8], [28], [49], [51], [65]. We furthermore incorporated clinical motor-cortex data from a large major-repression patient cohort (*N* = 4748).

The model is structured into three elements, specifically a spatial model, an orientation model, and an MEP recruitment model. A block diagram (Fig. 1) summarises how the location, orientation, and recruitment modules interact to predict the peak-to-peak MEP amplitude. All model elements are parametric and statistical so that the combination can describe and represent different subjects. Whereas the spatial and orientation model only contain inter-individuality, the MEP recruitment model further includes intra-individual stimulus-to-stimulus variability. The latter uses a previously developed MEP model [65]. The parameters of the elements represent an individual subject and can reproduce the same subject later again. The model furthermore offers a virtual subject generator that generates the parameters. It includes population statistics for the parameters to randomly generate a virtual subject population where no two subjects are alike.

**Fig. 1.**
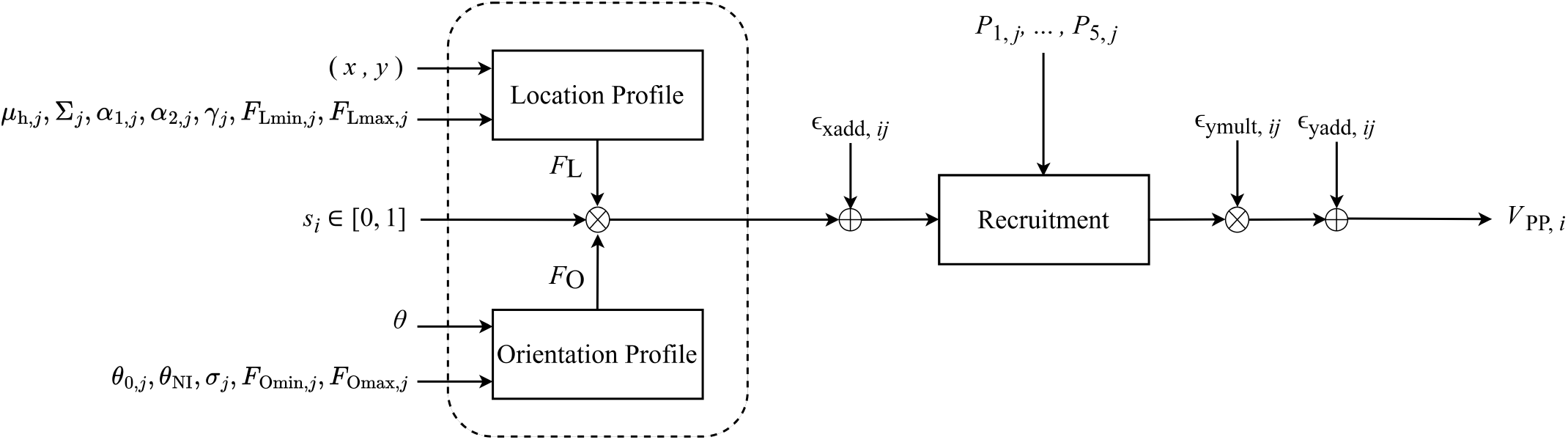
Block diagram of the overall location–orientation–recruitment model. The location and orientation components modulate the effective stimulation level, which the recruitment model converts into motor-evoked potentials (MEP) with a corresponding peak-to-peak amplitude. The input variables are *x, y, θ*, and *s*_*i*_. The intra-individual trial-to-trial variability disturbances *ϵ*_x add,*ij*_, *ϵ*_y add,*ij*_, and *ϵ*_y mult,*ij*_. The remaining variables represent the subject parametrically. On the one hand, those parameters identify and represent a specific individual subject. On the other hand, their statistical distribution (Figs. 3 and 5) represent the population.

### B. Spatial Model

#### 1) Population Level

We used our statistical data of the hotspot, i.e., the position with the typically largest MEP response for the population-level representation of the motor-hotspot location. The hotspot dataset was mapped into the surface coordinate system of the spatial model. The hotspot locations have a custom 1 cm resolution scalp grid over the left motor cortex, referenced to the vertex (Cz) [66]. The grid is anchored by a diagonal reference row defined by points located 5 cm anterior and 5 cm left of Cz. Each grid label was converted into a two-dimensional scalp coordinate. The resulting coordinates *µ*_h,*j*_ = (*x*_h,*j*_, *y*_h,*j*_)^*T*^ were then used to estimate the population mean ***µ*** and the covariance matrix **Σ** of the hotspot location per

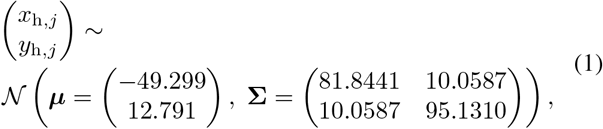

from which subject-specific hotspot centers are generated for the virtual-subject population. The subject-specific location *µ*_h,*j*_ = [*x*_h,*j*_, *y*_h,*j*_]^*T*^ denotes the maximum of the parametric individual spatial profile (see below). Hotspot locations were expressed in a Cz-centered scalp coordinate system.

Figure 2 illustrates a representative measured motor-map area derived from one participant in one session and overlaid on the three-dimensional MNI152 standard brain template, where yellow indicates the maximum MEP amplitude.

**Fig. 2.**
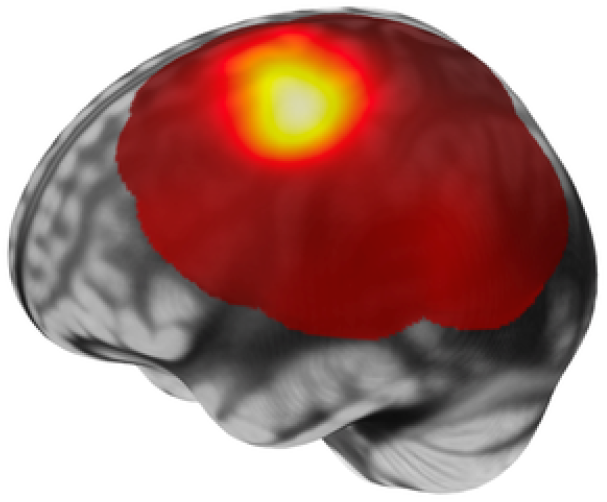
Representative measured motor-map area from one participant in one session overlaid on the three-dimensional MNI 152 standard brain template.

#### 2) Individual Spatial Profile

Although previously suggested, a multivariate normal function did not well represent the spatial map of MEP excitability within a subject, particularly due to its symmetry and inability to describe skewness of the profile. Due to the influence of gyrification, such incompatibility of a symmetric function is expected. We therefore implemented a *multivariate extended skew normal distribution* [67]. This function represents the spatial model, which provides the effective stimulation strength in the target location for a specific spatial coordinate.

Let **r** = (*x, y*)^*T*^ denote the two-dimensional spatial surface coordinate. The extended skew normal spatial kernel was defined as

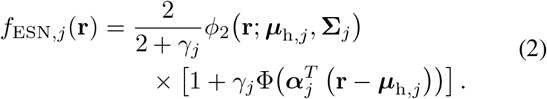

The location factor was obtained from the peak-normalized kernel as

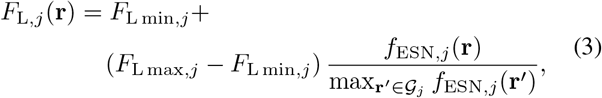

where *G*_*j*_ denotes the spatial search grid around ***µ***_h,*j*_ . *F*_L,*j*_ represents the location factor that provides the effective stimulus gain for subject *j* at a specific spatial surface coordinate **r** = (*x, y*)^*T*^, *F*_Lmin,*j*_ is the baseline location factor, and *F*_Lmax,*j*_ is the maximum location factor. *ϕ*_2_(**r**; ***µ***_h,*j*_, **Σ**_*j*_) is the two-dimensional normal density function with the mean vector *µ*_h,*j*_ and covariance matrix **Σ**_*j*_. Φ( ) denotes the cumulative gaussian function, *γ*_*j*_ the extended parameter, and ***α***_*j*_ = [*α*_1,*j*_ *α*_2,*j*_]^*T*^ the skewness parameter vector. Parameter *α*_1,*j*_ refers to the x-direction, and *α*_2,*j*_ the y-direction skewness.

The parameters of the spatial profile were also estimated from the experimental mapping data. We performed Shapiro– Wilk tests (values in parentheses report the *p*-value and test statistic *W* , all *p >* 0.05) to identify the population statistics of the parameters, which by themselves describe the individual spatial excitability profile with extended skew normal shape.

The population of our dataset demonstrated normal distributions with the parameters *α*_1,*j*_ (*p* = 0.983, *W* = 0.984) and *α*_2,*j*_ (*p* = 0.964, *W* = 0.982). Similarly, parameters *γ*_*j*_ (*p* = 0.414, *W* = 0.951), *F*_Lmax,*j*_ (*p* = 0.155, *W* = 0.924), and *F*_Lmin,*j*_ (*p* = 0.645, *W* = 0.964) were normally distributed. Parameter Σ_11,*j*_ (*p* = 0.2658, *W* = 0.9444) followed a normal distribution, Σ_12,*j*_ and Σ_22,*j*_ lognormal distributions so that lg(Σ_12,*j*_) *≡*log_10_(Σ_12,*j*_) (*p* = 0.4746, *W* = 0.9579) and lg(Σ_22,*j*_) (*p* = 0.3152, *W* = 0.9482) turned out normal.

Each parameter was represented as a distribution with its respective mean (*µ*) and variance (*σ*^2^) as

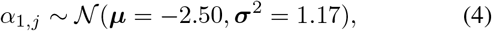

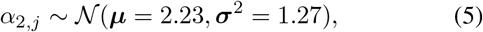

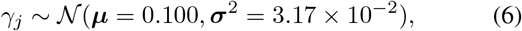

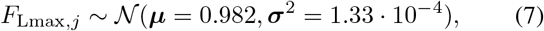

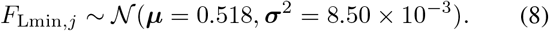

The covariance matrix Σ_*j*_ in Eq. (2) describes the spatial spread of the profile around that hotspot center. Its unique elements were modeled across subjects as

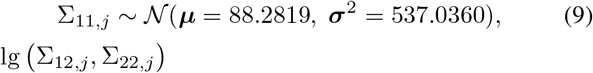

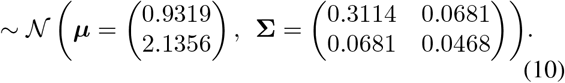

Figure 3 illustrates the found distribution of the location parameters in the population.

**Fig. 3.**
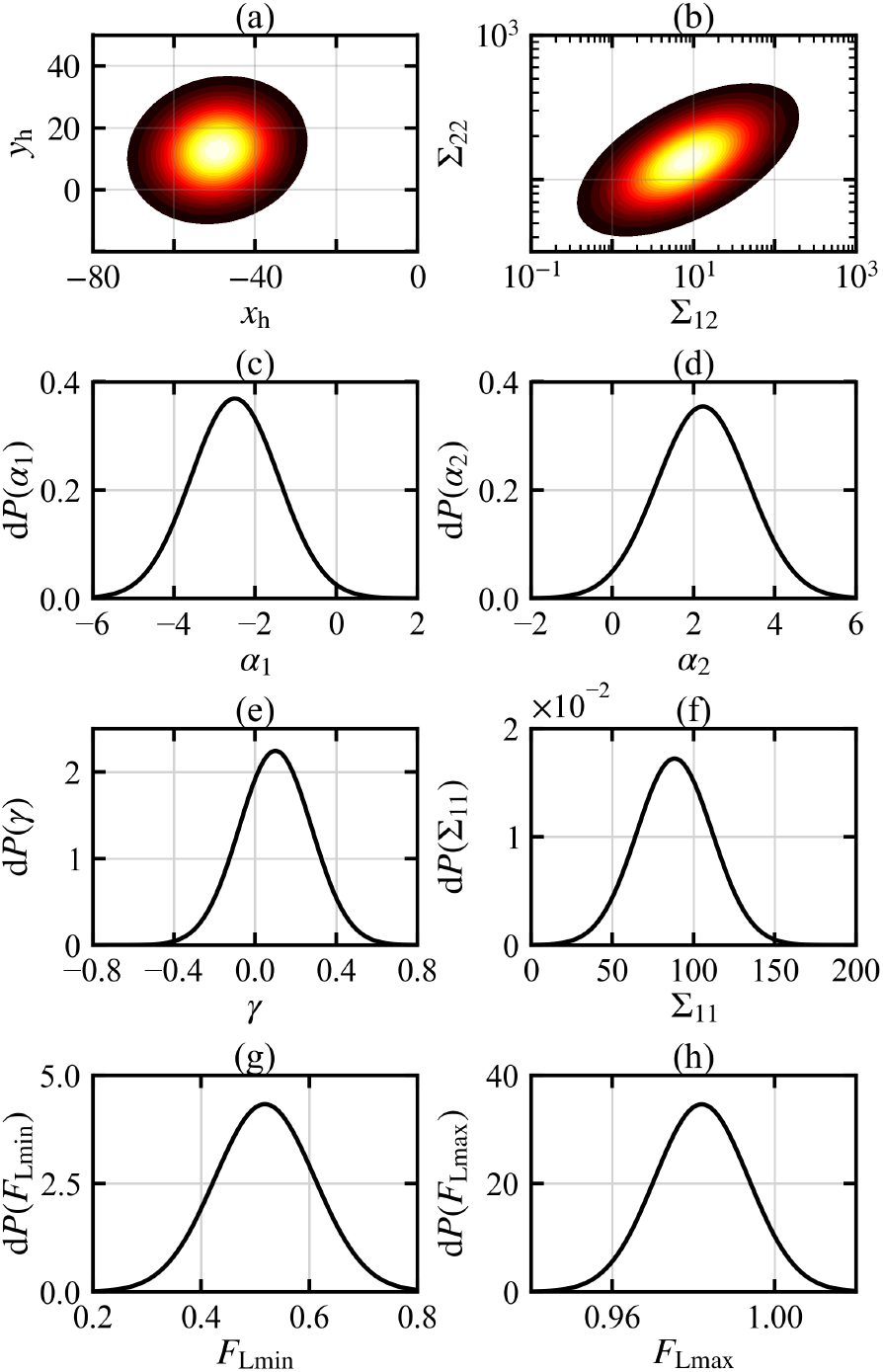
Statistical distributions of the model parameters in our population for the location model.

### C. Coil Orientation Modeling

The orientation model complements the location model with the impact of coil orientation, i.e., rotation around the surface normal, on the recruitment. The tangential alignment of the coil with the focus point on the local head surface is a question of operator skills rather than a practical degree of freedom in the use of TMS and was studied and modelled elsewhere [34], [35].

The orientation model represents the orientation factor that determines the effective stimulus gain as a function of coil orientation. The maximum of the curve typically occurs for coil orientations which generate an induced electric field approximately perpendicular to the central sulcus [33]. In this model, the orientation is expressed relative to the nasion–inion line, which serves as a consistent reference based on the 10–20 coordinate system. The angle between the nasion–inion line and the central sulcus varies and depends on the individual anatomy. Neither previously used cosine [68] nor Gaussian functions [69] provided a good fit to the coil orientation curve.

Figure 4 contrasts the generalised-logistic orientation model with Gaussian and cosine alternatives. We identified a generalized logistic function as more appropriate, which was previously suggested in the literature [70] and describes well the relationship between stimulus strength (input) and the neuronal output. However, a logistic function is monotonic and is therefore not directly suitable for describing the peak-shaped orientation dependence. To overcome this limitation, we split the orientation curve into monotonic angular portions around the posterior–anterior (PA) and anterior–posterior (AP) directions. In this way, the PA and AP portions of the curve could have individual parameters to reflect their different shape and excitability [33]. The generic logistic form for each quadrant is

**Fig. 4.**
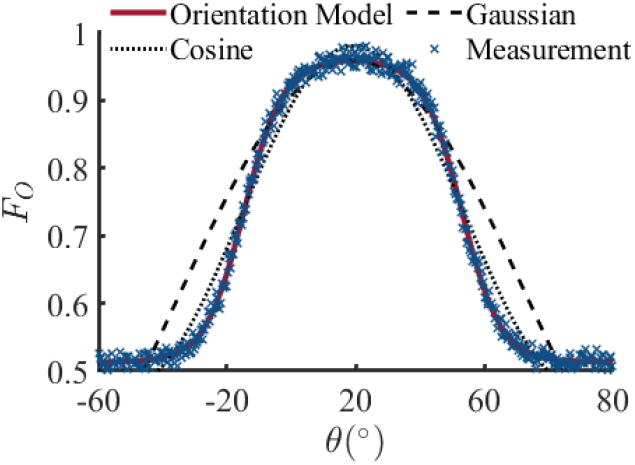
Modeling of the orientation factor as a function of coil / field orientation. Comparison of the logistic equation (solid line), cosine function (dotted line), and Gaussian function (dashed line).

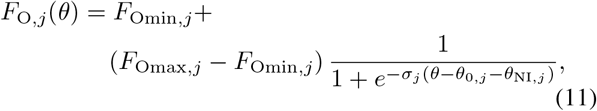

where *F*_O,*j*_ represents the orientation factor for subject *j* as a function of coil orientation *θ*. The terms *F*_Omin,*j*_ and *F*_Omax,*j*_ respectively denote the minimum and maximum orientation factors. *θ*_0,*j*_ is the center of the logistic response curve, *θ*_NI,*j*_ is the angle between the line perpendicular to the central sulcus and the nasion–inion reference line, and *σ*_*j*_ is the slope of the logistic curve.

We estimated the parameters separately for AP as well as PA orientations and tested for normality using the Shapiro–Wilk test. The parameters *θ*_0,PA,*j*_ (*p* = 0.930, *W* = 0.977), *θ*_0,AP,*j*_ (*p* = 0.546, *W* = 0.953), *σ*_PA,*j*_ (*p* = 0.803, *W* = 0.968) and *σ*_AP,*j*_ (*p* = 0.451, *W* = 0.948) suggest normal distributions. Similarly, *F*_Omax,PA,*j*_ (*p* = 0.105, *W* = 0.921), *F*_Omax,AP,*j*_ (*p* = 0.139, *W* = 0.928) and *F*_Omin,PA,*j*_ (*p* = 0.347, *W* = 0.949) also follow normal distributions.

Previous measured coil orientation maxima in the literature do not necessarily agree with each other (*p <* 0.01) [71], [72]. Distributions rarely overlap and the difference between means is larger than, sometimes multiple times, their estimated standard intra-study deviation. Thus, individual studies appear to be under-powered and to under-estimate the real population spread. We modelled each parameter with individual normal distribution (mean *µ* and variance *σ*^2^) as

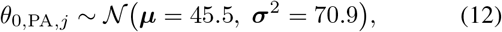

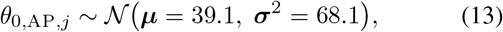

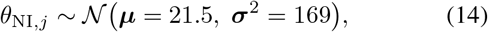

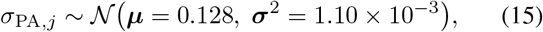

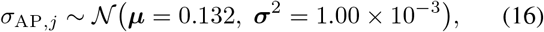

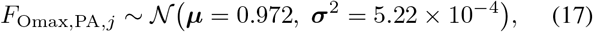

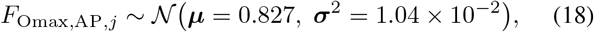

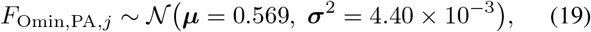

and *F*_Omin,AP,*j*_ is obtained by continuity.

Figure 5 illustrates the found distribution of the orientation parameters in the population.

**Fig. 5.**
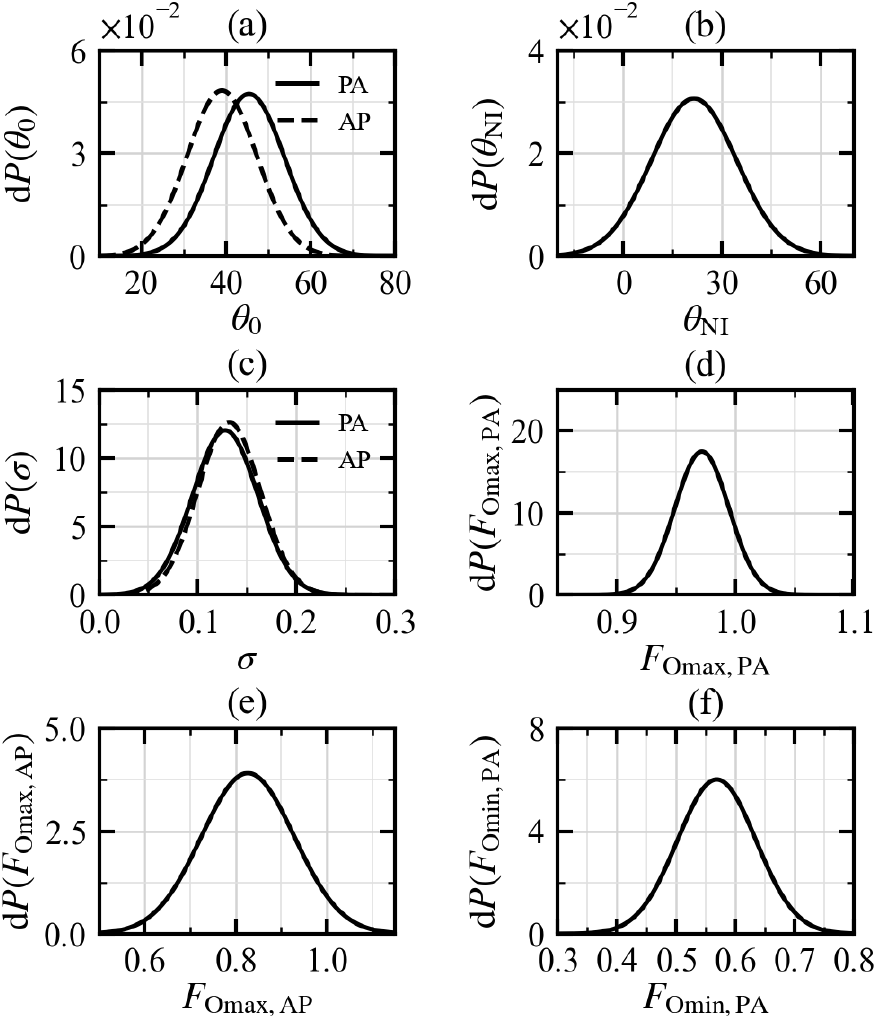
Statistical distributions of the parameters of the generalized logistic equation.

### D. Integration of Orientation and Location Models into the Recruitment Framework

The fundamental recruitment model outputs the peak-to-peak MEP voltage and follows

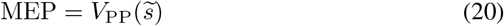

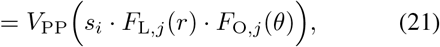

which combines the spatial (*x, y*) and orientational (*θ*) mod-els with the recruitment ^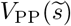^ of the effective stimulation strength 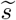 at the target per

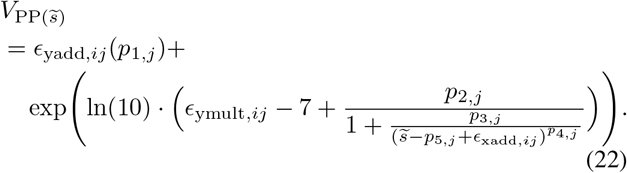

The recruitment model was adopted from the literature [65]. Stimulation strength *s* represents the stimulator output, e.g., 0 … 1 maximum stimulator output. In this model, the parameters *p*_1,*j*_, … , *p*_5,*j*_ are the individual recruitment parameters for subject *j*. The terms *ϵ*_*x*add,*ij*_, *ϵ*_*y*mult,*ij*_, and *ϵ*_*y*add,*ij*_ represent trial-to-trial variability terms added to the recruitment model. Specifically, *ϵ*_*x*add,*ij*_ is an additive input-side variability term, *ϵ*_*y*mult,*ij*_ a multiplicative output-side variability term, and *ϵ*_*y*add,*ij*_ an additive output-side physiological/measurement-noise term. The statistical distributions of the recruitment parameters and variability terms were taken from the original statistical MEP model [65].

## III. Combined Model Behavior

The model allows the generation of virtual subjects of a population whose individual parameters are sampled from the above distributions. A set of parameters represents a specific subject with the inter-individual spread of parameters. Sampling from the parameter distributions generates a random new individual. In addition to subject individuality, the responses of a subject furthermore demonstrate the intra-individual variability with the detailed features known from the literature [13], [22], [65], [73], [74].

The model allows large-scale testing of novel closed-loop methods at the design stage with very diverse subjects that represent a larger population. The numbers of subjects and test pulses can be several powers of ten larger than a single experimental study may ever allow. Furthermore, the model avoids safety problems of early-stage closed-loop procedures, which can demonstrate erratic behaviour and should not be run on humans initially without such prior large-scale *in-silico* testing.

Figure 6 visualises the position–orientation–recruitment model with several for five random subjects. The graphs show several location and recruitment curves of different virtual subjects with their intra-individual trail-to-trial variability. The parameters were randomly selected based on the identified distributions to represent a random individual from the population (subject constructors to sample a random subject using population statistics part of model and code, Figs. 3 and 5).

**Fig. 6.**
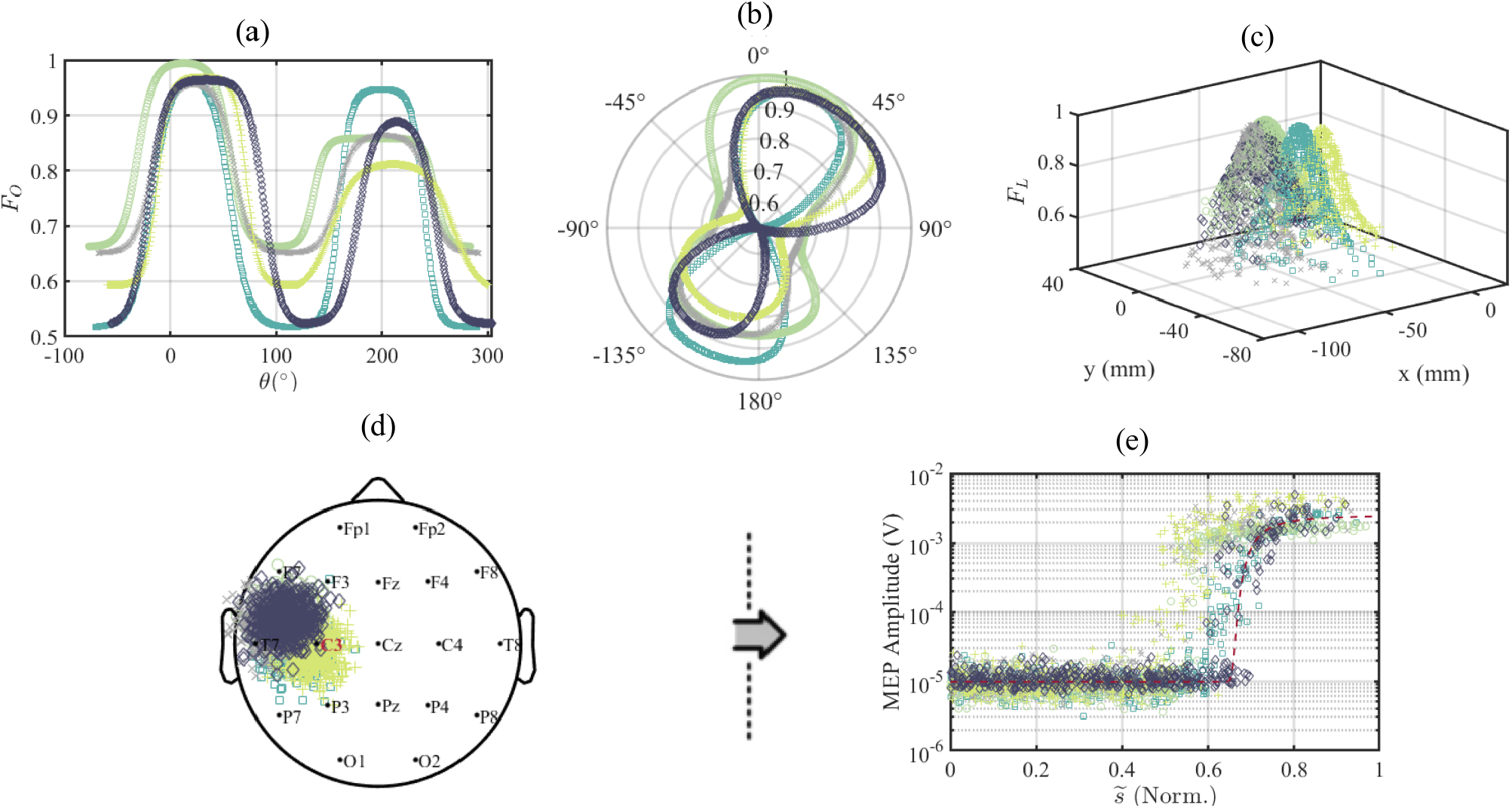
Visualisation of typical output of the model for five randomly sampled subjects. (a, b) Orientation factor *F*_O_ as a function of coil orientation *θ* for the five representative virtual subjects. (c) Location factor *F*_L_ over scalp coordinates (*x, y*), which demonstrates inter-individual differences in hotspot location and profile. (d) Projection of measured samples onto the 10–20 scalp coordinate system. (e) Corresponding recruitment input–output clouds with both inter-individual differences in recruitment behavior and intra-individual trial-to-trial variability.

The spatial distribution of the location model was visualized within the 10–20 EEG coordinate system, constructed based on a combination of cranial landmarks (nasion, inion, left and right pre-auricular points). For visualization purposes, the spatial map was projected onto a standardized scalp representation. The population-level hotspot as the maximum of the excitability profile landed close to the C3 position, which is known to be a good first estimate for the typical hand representation of the left hemisphere. The spatial maps demonstrate skewness, which was learned from the experimental data and represents the influence of the cortical gyrification [75]. The orientation leads to two peaks. The one in posterior–anterior direction is higher and not far from 45°, which reflects reports from the literature [33]. The anterior–posterior peak is smaller and by approximately 180° shifted.

The recruitment demonstrates typical sigmoidal behavior with a baseline formed by background activity and noise and a characteristic skewed extreme-value distribution for low stimulation strength, a rising slope in which the responses rapidly increase on average for stronger stimuli but with high trial-to-trial variability due to an interaction of excitability fluctuations as well as output variability, and a saturation plateau in which the mostly log-normal variability decreases again. Similar to the shape parameters, also the intra-individual variability properties are sampled from statistical distributions and vary from subject to subject.

## IV. Conclusion

This paper introduced an integrated model to support development and test of closed-loop TMS methods on virtual subjects in a software-in-the-loop fashion. The model includes a spatial (cortical coil location and orientation) and recruitment component to generate a matching MEP amplitude for a specific pulse strength administered to a certain location with a given coil orientation. The model was calibrated to the population represented by study data and represents inter-subject differences in spatial and recruitment behavior. Furthermore, the model also includes intra-individual variability and integrates location and orientation into the recruitment framework. The model successfully simulates virtual subjects with a wide range of characteristics to reflect anatomical and physiologic diversity, which is important for the development and tuning of better (closed-loop) methods.

## V. Code Availability

An implementation of the model in MATLAB accompanied by instructions for its use is available at https://github.com/mfarahmandrad/Spatial-Recruitment-Model. This model can be freely used and improved by the research community.

## IV. Acknowledgment

The authors want to thank Brett Cormier and Katherine Kriske of Salience Health for their willingness to share techniques and data regarding hotspot localization.

